# Blockade of ghrelin receptor signaling enhances conditioned passive avoidance and context-associated neural circuit activation in fasted male rats

**DOI:** 10.1101/2022.02.11.480168

**Authors:** Caitlyn M. Edwards, Inge Estefania Guerrero, Tyla Dolezel, Huiyuan Zheng, Linda Rinaman

**Author notes:** Corresponding Author: Linda Rinaman, Ph.D., Department of Psychology, Florida State University, 1107 W. Call Street, Tallahassee, FL, 32304, USA.

## Abstract

Interoceptive feedback to the brain regarding the body’s physiological state plays an important role in guiding motivated behaviors. For example, a state of negative energy balance tends to increase exploratory/food-seeking behaviors while reducing avoidance behaviors. We recently reported that overnight food deprivation reduces conditioned passive avoidance behavior in male (but not female) rats. Since fasting increases circulating levels of ghrelin, we hypothesized that ghrelin signaling contributes to the ability of fasting to reduce conditioned avoidance. To test this, *ad libitum*-fed male rats were trained in a passive avoidance procedure using mild footshock. Later, following overnight food deprivation, the same rats were pretreated with ghrelin receptor antagonist (GRA) or saline vehicle 30 min before avoidance testing. GRA restored passive avoidance in fasted rats as measured both by latency to enter and time spent in the shock-paired context. In addition, compared to vehicle-injected fasted rats, fasted rats that received GRA before re-exposure to the shock-paired context displayed more activation of prolactin-releasing peptide (PrRP)-positive noradrenergic neurons in the caudal nucleus of the solitary tract, accompanied by more activation of downstream neurons in the bed nucleus of the stria terminalis and paraventricular nucleus of the hypothalamus. These results support the view that ghrelin signaling contributes to the inhibitory effect of fasting on learned passive avoidance behavior, perhaps by suppressing recruitment of PrRP-positive noradrenergic neurons and their downstream hypothalamic and limbic forebrain targets.

## Introduction

Avoiding danger is critical for survival, and animals are equipped with neural circuits that enable adaptive avoidance responses to stimuli previously associated with potentially dangerous, aversive experiences. However, excessive conditioned avoidance can be maladaptive, contributing to clinical anxiety and other stress-related disorders [1–3]. Conditioned avoidance responses are based on learning and memory processes that include both conscious and subconscious components. Sensory feedback from body to brain during the learning and expression of emotional responses plays a major role in the subconscious modulation of motivated behavior [4]. For example, short-term negative energy balance generally biases animals towards approach behaviors and away from avoidance behaviors, thereby facilitating exploratory food-seeking [5,6]. In recent work, we demonstrated a reduction in learned passive avoidance behavior in male rats that were tested following an overnight fast [7]. The present study was designed to investigate potential neural circuits through which overnight food deprivation suppresses conditioned passive avoidance behavior.

Interoceptive feedback about the internal state of the body is continuously communicated to the brain through hormonal and neural pathways. The neural pathways feature vagal afferent inputs to the caudal nucleus of the solitary tract (cNTS) that can powerfully modulate motivated behavior [4,8–10]. Vagal afferents communicate a diverse array of cardiovascular, respiratory, gastrointestinal, and inflammatory signals to the cNTS in a manner that is further modulated by hormonal and metabolic state [11–15]. Within the cNTS, the A2 noradrenergic cell group receives direct synaptic input from vagal sensory afferents [16,17], and A2 neurons are activated to express the immediate-early gene product, cFos, in response to vagal sensory stimulation [17–20]. A caudal subset of A2 neurons that co-express prolactin-releasing peptide (PrRP) express cFos in rats exposed to a variety of innate and conditioned stressors that promote avoidance behavior [6,7,21,22]. Interestingly, overnight food deprivation markedly reduces the ability of these innate and conditioned stress stimuli to activate PrRP+ neurons within the cNTS [6,7], concurrent with reduced activation of neurons within hypothalamic and limbic forebrain regions that are innervated by PrRP+ neurons, including the paraventricular nucleus of the hypothalamus (PVN) [19] and the anterior ventrolateral bed nucleus of the stria terminalis (vlBNST) [6,7].

Food deprivation increases circulating levels of acyl-ghrelin [23–25], and acyl-ghrelin reduces tonic firing and stimulus-induced responsiveness of vagal sensory neurons [26,27] and postsynaptic A2 neurons in the cNTS [28]. We recently demonstrated that systemic administration of a ghrelin receptor antagonist in fasted rats partially restores the ability of vagal sensory stimulation to activate PrRP+ A2 neurons and suppress food intake [20]. Further, caloric restriction reduces innate avoidance behavior in wildtype mice but not in mice lacking ghrelin receptors [5], which display more innate anxiety-like behavior than wildtype mice [29]. Consistent with these anxiolytic effects of ghrelin signaling, baseline acyl-ghrelin levels prior to auditory fear conditioning negatively predict conditioned freezing behavior in rats [30]. However, the potential impact of ghrelin signaling on the expression of learned avoidance has not been examined. Based on our previous research demonstrating that rats fasted overnight display less passive avoidance behavior and less fear context-induced neural activation [7], the present study was designed to test the hypothesis that increased ghrelin receptor signaling contributes to the ability of overnight food deprivation to suppress learned passive avoidance behavior, and to suppress conditioned context-induced activation of PrRP+ neurons in the cNTS and their downstream targets in the PVN and anterior vlBNST.

## Materials and Methods

All experiments were conducted in accordance with the National Institutes of Health *Guide for the Care and Use of Laboratory Animals* and were reviewed and approved by the Florida State University Animal Care and Use Committee.

### Passive Avoidance Task

Adult male Sprague Dawley rats [Envigo; N=32 (250-300g body weight)] were pair-housed in standard tub cages in a temperature- and light-controlled housing environment (lights on from 0400 hr to 1600 hr). Only male rats were used in the current study based on evidence that food deprivation does not significantly reduce conditioned passive avoidance in female rats [7]. Rats were acclimated to handling for three days, with free access to water and rat chow (Purina 5001). Rats underwent passive avoidance training in a novel light/dark shuttle box (Colbourn Instruments, Allentown, PA), with training performed 4-6 hr after lights on. The light/dark box comprised a light (illuminated) chamber with clear plastic walls and a smooth white plastic floor, and a dark (non-illuminated) chamber with black plastic walls and a metal grid floor. Each chamber measured 25 × 25 cm, with 28 cm-high walls and a ceiling. The two chambers were separated by a metal divider wall with a guillotine door that could be opened and closed remotely. For passive avoidance training, rats were initially placed individually into the light chamber of the box and the dividing guillotine door was lifted immediately to allow access to the dark chamber. As expected, due to their innate aversion to the light and preference for the dark, rats very quickly entered the dark chamber (average latency = 22.73s). Upon entry into the dark chamber, the guillotine door was closed. After a 5s delay, rats received a single mild electric footshock (0.6mA; 1s). Rats remained in the enclosed dark chamber for 30s following footshock, and then were returned to their homecage. Cagemates were similarly trained on the same day.

Two hours before dark onset on the day before testing (i.e., 2 days after training), all rats were transferred to clean cages with chow removed to initiate overnight food deprivation. On the next day, 4-6 hr after lights on, rats received either an intraperitoneal (i.p.) injection of 0.15M NaCl (saline) vehicle (∼0.3 ml), or the same volume of vehicle containing ghrelin receptor antagonist (GRA; [D-Lys3]-GHRP-6; 3.3mg/kg BW; Sigma-Aldrich G4535). This GRA dose was based on our previous report demonstrating its ability to restore cholecystokinin-induced NTS neuronal activation in food deprived rats [20]. Thirty minutes later, rats were tested for passive avoidance retention. For this purpose, rats were placed individually into the light chamber of the light/dark box, the guillotine door was opened, and the latency of rats to fully enter the dark chamber was recorded, with a preset maximum latency of 900s (15 min). During the retention test, the guillotine door remained open and no footshock was administered. Rats were allowed to freely explore both the light and dark chambers during the 900s test, and total time spent within each chamber was recorded. After testing was complete, rats were returned to their home cages. Chow was not returned until 2 hr prior to dark onset (i.e., approximately 6-8 hr after testing).

To confirm that GRA treatment does not alter conditioned avoidance behavior in non-fasted rats, a separate cohort of adult male rats (n=4) was trained in the passive avoidance task as described above. Two days later, rats were tested in an *ad libitum* fed state, 30 min after receiving i.p. injection of GRA.

### Novel Object Recognition Task

To examine the impact of GRA on general memory function, rats (N=32) were tested for novel object recognition at least 5 days after completing passive avoidance testing. For this, rats underwent 3 days of acclimation (10 min/day) to a previously novel tub cage that had a thin layer of bedding on the floor, with the tub enclosed in an illuminated sound-attenuating chamber. On the 4th day, rats underwent object training 4-6 hr after lights on, during which two similarly sized but geometrically distinct ceramic objects with rounded contours (i.e., a white pyramidal “Eiffel tower” and a blue spherical “rocket ship”) were placed at opposite ends of the tub cage. Rats were allowed to freely explore both objects for 15 minutes while their behavior was recorded and later analyzed using behavior analysis software (ANY-maze, Stoelting Co.). During training, there was no significant difference between time spent with either object, defined as the rat being in contact with or actively examining the object.

Two hours before dark onset on the day before testing (i.e., 2 days after object training), all rats were transferred to clean cages with chow removed. On the subsequent day, fasted rats received either an i.p. injection of saline vehicle (∼0.3 ml), or the same volume of vehicle containing GRA (3.3mg/kg BW). Thirty minutes later, rats were tested for novel object recognition. For this, the previously explored “Eiffel tower” object was replaced with a novel object (i.e., a grey cube-shaped “robot”). Rats were allowed to freely explore the novel and familiar objects for 15 minutes. The time each rat spent exploring each object was recorded, and the discrimination index was calculated as the difference between time spent with each object divided by the total time spent with either object:

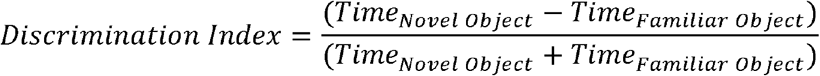

In the discrimination index, a value of 0 indicates no discrimination between the novel and familiar objects. A positive value indicates more time spent with the novel object, whereas a negative value indicates more time spent with the familiar object. One rat from each injection group was excluded from analysis after one of the objects was knocked over during the novel object recognition test.

### Open Field Test

To examine the impact of GRA on general locomotion and anxiety-like behavior, the same rats (N=32) were tested in the open field after overnight food deprivation. On the testing day (4-6 hr after lights on), fasted rats received either an i.p. injection of saline vehicle (∼0.3 ml), or the same volume of vehicle containing GRA (3.3mg/kg BW). Thirty minutes later, rats were placed into the center of a novel 95 × 95cm open arena, and then left to freely explore for 10 min. Behavior in the open field was recorded using a Logitech webcam and was analyzed using automated behavior analysis software (ANY-maze, Stoelting Co.). Locomotion was defined as the total distance traveled during the test. Anxiety-like behaviors were interpreted based on time spent in the center zone of the field (∼30 × 30 cm) and latency to first enter the center zone. Data from one cohort of rats (n=4/injection group) were excluded due to lighting issues in the testing room.

### Conditioned Context Re-exposure and Perfusion

To examine neural activation in response to the conditioned context associated with footshock, a subset of rats used in the behavioral studies described above (N=24) were retrained in the passive avoidance task, in which they were placed again into the light side of the light/dark box. When rats entered the dark chamber (average latency to enter = 134s, reflecting their earlier shock conditioning session), the guillotine door closed behind them. Shock-retrained rats immediately received a mild electric footshock, as they did during their initial conditioned avoidance training session. Additional shock-naive control rats (N = 12) were placed into the light side of the light/dark box but did not receive any footshock upon entry into the dark chamber (average latency to enter = 19s, similar to that in our previous report [7]). Rats in both the shock retrained (i.e., shocked) and non-shocked groups were kept in the dark chamber for 30s before being returned to their home cages. Two days later, 2 hr before dark onset, all rats were transferred to clean cages with chow removed to initiate overnight food deprivation. On the next day, 4-6 hr after lights on, fasted rats in both groups received either an i.p. injection of 0.15M NaCl (saline) vehicle (∼0.3 ml), or the same volume of vehicle containing GRA (3.3mg/kg BW). Thirty minutes later, rats were placed directly into the dark chamber, which was an aversively conditioned stimulus only in the shock-retrained group. Rats remained in the dark chamber for 10 min with the guillotine door closed, and were then returned to their home cages. Sixty minutes later, rats were deeply anesthetized with pentobarbital sodium (39 mg/ml i.p., Fatal Plus Solution; Butler Schein) and transcardially perfused with saline (100mL) followed by 4% paraformaldehyde (500mL).

### Histology

Fixed brains were removed from the skull, post-fixed overnight at 4°C, then cryoprotected in 20% sucrose. Brains were blocked and sectioned coronally (35 μm) using a Leica freezing-stage sliding microscope. Tissue sections were collected in six serial sets and stored at -20°C in cryopreservant solution [31] until immunohistochemical processing. Primary and secondary antisera were diluted in 0.1M phosphate buffer containing 0.3% Triton X-100 and 1% normal donkey serum. One set of tissue sections from each rat (with each set containing a complete rostrocaudal series of sections spaced by 210 μm) was incubated in a rabbit polyclonal antiserum against cFos (1:10,000; Cell Signaling, 2250; AB_2247211), followed by biotinylated donkey anti-rabbit IgG (1:500; Jackson ImmunoResearch). Sections were then treated with Elite Vectastain ABC reagents (Vector) and reacted with diaminobenzidine (DAB) intensified with nickel sulfate to produce a blue-black nuclear cFos reaction product.

To visualize cFos within hindbrain PrRP neurons, one set of cFos-labeled tissue sections from each rat was subsequently incubated in a rabbit polyclonal antiserum against PrRP (1:10,000; Phoenix Pharmaceuticals, H-008-52), followed by biotinylated donkey anti-rabbit IgG (1:500; Jackson ImmunoResearch) and Elite Vectastain ABC reagents (Vector), and finally reacted with plain DAB to produce a brown cytoplasmic reaction product.

To visualize cFos within all noradrenergic neurons, a second set of tissue from each rat was incubated in a cocktail of rabbit anti-cFos (1:1000) and a mouse monoclonal antiserum against dopamine beta-hydroxylase (DbH; 1:5000; Millipore, MAB308; AB_2245740), followed by a cocktail of Cy3-conjugated donkey anti-rabbit IgG (red) and Alexa Fluor 488-conjugated donkey anti-mouse IgG (green).

### Quantification of Neural Activation

Sections through the cNTS (∼15.46-13.15 mm caudal to bregma) were imaged at 20x using a Keyence microscope (BZ-X700). Using separate sets of dual-labeled tissue sections from each rat, the total numbers of DbH+ and PrRP+ cNTS neurons were counted, and the percentage of each chemically identified neural population co-expressing cFos was determined. A neuron was considered to be cFos-positive if its nucleus contained any visible cFos immunolabeling.

Sections through the anterior vlBNST (∼0.26-0.40mm caudal to bregma) were imaged at 10x. Images from two rats were excluded from further analysis due to poor tissue quality. Using ImageJ, a circular region of interest (ROI) of 0.10mm^2^ diameter was centered on the area containing the highest density of DbH+ fibers and terminals, corresponding to the fusiform subnucleus of the anterior vlBNST [6,32,33]. The number of cFos-positive neurons within each 0.10mm^2^ ROI was determined bilaterally in 2-3 sections per brain and then averaged across assessed ROIs in each animal.

Three sections through the PVN (∼1.5-1.9mm caudal to bregma) in each animal were imaged at 10x. For this, a region of interest (ROI) encompassing each PVN (bilaterally) was identified by the presence of dense DbH+ fibers and terminals, and an outline was drawn around each region using ImageJ. The number of cFos-positive neurons within each ROI was determined, divided by the ROI area, and averaged in each animal to calculate the number of cFos-positive neurons per mm^2^.

### Ghrelin ELISA

After finding that GRA treatment significantly increased conditioned avoidance behavior and cFos activation in fasted rats (see *Results*), we sought to confirm our assumption that plasma acyl-ghrelin levels increase in rats exposed to either a single or repeated sessions of overnight food deprivation. For this, terminal blood samples were collected from a separate cohort of adult male rats that were either *ad lib* fed with no fasting experience (n=3), fasted once overnight before morning blood collection (n=3), or fasted overnight on 3 separate occasions (with at least two days between each episode) with the 3rd fast occurring the night before morning blood collection (n=4). Just before blood collection, rats were injected with pentobarbital sodium (39 mg/ml i.p., Fatal Plus Solution; Butler Schein), and blood samples were collected within 5 min from the right atrium of the heart using a 3mL syringe pre-rinsed with 1% EDTA. Blood samples were transferred to a K2E EDTA vacuette tube (BD Ref 367841), and aprotinin (50µl/mL) was added to each sample. Samples were centrifuged at 3500 rpm for 10 min at 4°C to separate the plasma. Hydrochloride was added to the plasma supernatant (10µl/100mL), and then samples were centrifuged again at 3500 rpm for 5 min at 4°C. Plasma samples were stored at -80°C until acyl-ghrelin was measured using an ELISA kit (BioVendor, R&D; RA394062400R), following the manufacturer’s instructions.

### Statistics

Data were analyzed using GraphPad Prism. Passive avoidance behaviors (latency to enter and total time spent in the dark chamber) during the retention test were analyzed using a Welch’s t-test [34]. Time spent exploring novel vs. familiar objects in the novel object recognition task was analyzed with a mixed 2×2 ANOVA with i.p. injection as a between-subjects independent variable and Object (novel, familiar) as a within-subjects independent variable. When ANOVA F values indicated significant main effects and/or interactions, post-hoc comparisons were made using Šídák’s multiple comparisons tests. Discrimination indices for novel object recognition by each i.p. injection group of rats were analyzed using Welch’s t-test, and by individual one-subject t-tests with a hypothetical value of 0 (i.e., no discrimination). Each open field behavioral measure (distance traveled, time in center, and latency to first enter the center) was analyzed separately with a Welch’s t-test to compare i.p. injection groups. Cell count data (cFos activation of PrRP and DbH neurons, cFos activation within the anterior vlBNST and PVN) were analyzed using between-subjects 2×2 ANOVA with previous shock-pairing and i.p. treatment (i.e., GRA vs. vehicle) as independent variables. In addition, Pearson’s r correlations were run between the percent of DbH+/PrRP+ cNTS neurons expressing cFos and cFos counts within vlBNST/PVN in the same rat. Plasma acyl-ghrelin data were analyzed using a one-way ANOVA with food deprivation history as the independent variable. An alpha level of 0.05 (p ≤ 0.05) was used as the criterion for considering group differences to be statistically significant. Estimation statistics [35,36] were used to report effect sizes for both passive avoidance behavior and cell count data.

## Results

### Passive Avoidance Task

Welch’s t-test indicated that fasted rats injected with GRA demonstrated more passive avoidance behavior than fasted rats injected with saline. Compared to saline-treated rats, GRA-treated rats took longer to enter the shock-paired dark chamber [t(26.1) = 1.728; p = 0.0479] and spent less time in the dark chamber [t(18.67) = 2.162; p = 0.0219] during the 900s testing period (Fig. 1). Estimation statistics indicated that compared to fasted saline-treated control rats, fasted GRA-treated rats took an average of 195s (3.25min) longer to enter the shock-paired dark chamber [95%CI: -11.4s, 419s], and spent an average of 178s (2.97min) less time in the shock-paired dark chamber during the 900s retention test [95%CI: 47.2s, 369s], evidence for increased passive avoidance after GRA treatment.

**Fig. 1.**
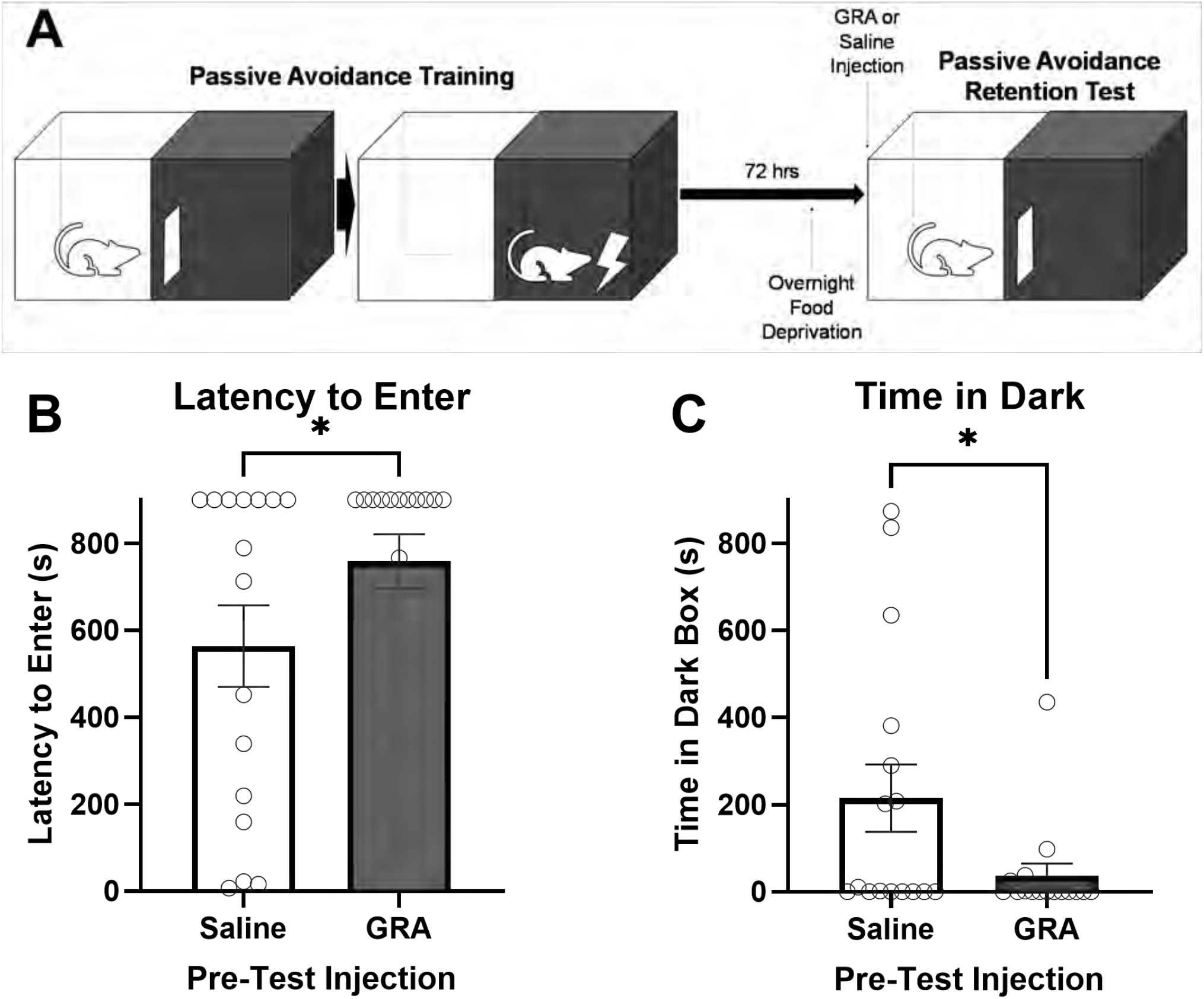
Systemic ghrelin receptor antagonist (GRA) administration prior to the retention test increased passive avoidance behavior in food deprived male rats. (A) Schematic depicting the passive avoidance experimental timeline. (B) Rats pretreated with GRA took an average of 195s longer to enter the shock-paired dark chamber compared to rats pretreated with saline. (C) Rats pretreated with GRA spent an average of 178s less in the shock-paired dark chamber during the 900s retention test compared to rats pretreated with saline. *p < 0.05. Graphs display group mean +/-SEM, with individual data points plotted.

A separate group of avoidance-trained, non-fasted rats that received GRA treatment before the passive avoidance test (n=4) displayed a typically long average latency to enter [895.75s ± 4.25] and minimal time spent within the shock-paired dark chamber [1s ± 1], quite similar to the behavior of fasted GRA-treated rats (see Fig. 1). The similarly robust conditioned avoidance behavior displayed by GRA-treated rats tested in the fasted and fed states also was similar to published data from our laboratory obtained from avoidance-trained, non-fasted male rats with no i.p. treatment prior to testing [7]. These observations support the view that GRA treatment does not alter conditioned avoidance behavior in *ad libitum-*fed rats, in which plasma acyl-ghrelin levels are not elevated.

### Novel Object Recognition Task

GRA treatment did not alter the behavior of fasted rats in the novel object recognition task (Fig. 2). Two-way ANOVA indicated a significant main effect of Object [F(1,28) = 15.59; p = 0.0005] but no main effect of i.p. treatment and no interaction. Post-hoc Šídák’s multiple comparisons tests indicated that fasted rats spent significantly more time exploring the novel object compared to the familiar object in both the saline-(p = 0.0093) and GRA-injected (p = 0.0361) groups. Individual one-sample t-tests indicated that the discrimination indices for both the saline-[t(14) = 3.575, p = 0.0030] and GRA-injected groups [t(14) = 2.294, p = 0.0378] were significantly higher than 0, evidence that both groups were similarly able to discriminate between the familiar and novel objects.

**Fig. 2.**
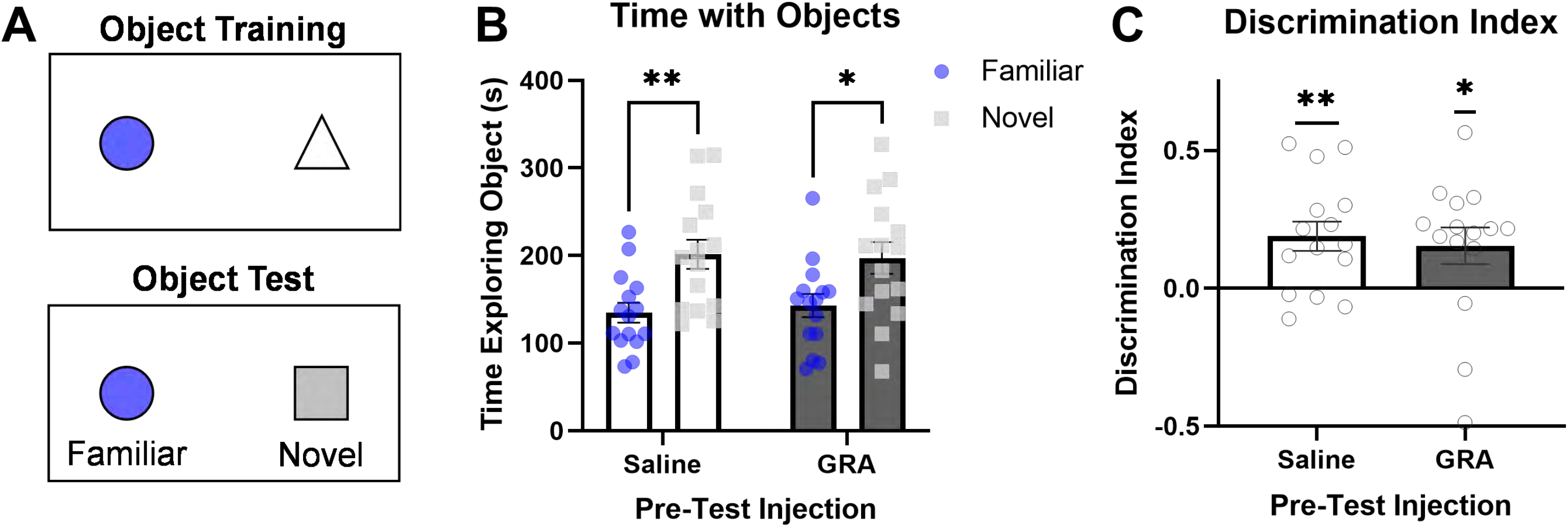
Systemic ghrelin receptor antagonist (GRA) administration prior to the novel object test did not influence novel object recognition memory in food deprived male rats. (A) Schematic depicting the novel object recognition training and test. (B) Both GRA- and saline-pretreated rats spent significantly more time with the novel object compared to the familiar object. (C) The discrimination indices for GRA- and saline-pretreated rats were similar, and significantly greater than zero in both groups. Graphs display group mean +/-SEM, with individual data points plotted. *p<0.05 and **p<0.01.

### Open Field Test

Pre-treatment of fasted rats with GRA did not alter assessed behaviors in the 10-min open field test (Fig. 3). There were no significant differences in the total distance traveled by saline-injected (mean = 17.9m) vs. GRA-injected rats (mean = 17.3m; Fig. 3A). Additionally, there were no significant group differences in anxiety-like behaviors as measured by either time spend in the central zone (Fig. 3B) or latency to first enter the central zone of the open field (Fig. 3C).

**Fig. 3.**
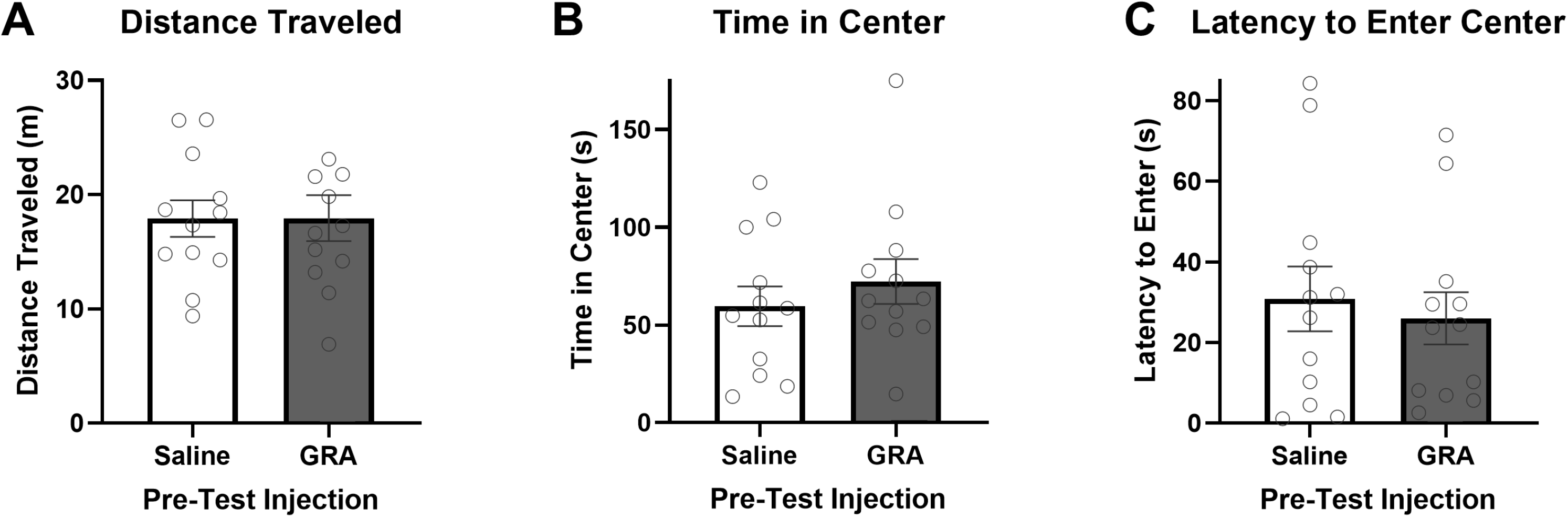
Systemic ghrelin receptor antagonist (GRA) administration prior to the open field test did not influence locomotion or anxiety-like behavior in food deprived male rats. (A) GRA- and saline-pretreated rats did not differ in distance traveled during the open field test. (B) GRA- and saline-pretreated rats did not differ in time spent in the center zone of the open field. (C) GRA- and saline-pretreated rats did not differ in latency to enter the center zone of the open field during the test. Graphs display group mean +/-SEM, with individual data points plotted.

### Neural Activation after Dark Chamber Re-exposure

Two-way ANOVA revealed a significant main effect of shock retraining [F(1,30 = 14.02; p < 0.001] on PrRP+ neuron activation in fasted rats that were re-exposed to the dark (i.e., shock-paired) chamber, as well as a trending main effect of GRA treatment prior to dark chamber re-exposure [F(1,30) = 3.14; p = 0.087] and a trending interaction between shock retraining and GRA treatment [F(1,30) = 3.33; p = 0.078] (Fig. 4A). Post-hoc Sidak’s multiple comparisons tests indicated that significantly more PrRP+ neurons were activated in shock-retrained fasted rats treated with GRA vs. rats treated with saline before dark chamber re-exposure (p = 0.010). Conversely, there was no GRA-related effect on the very low levels of PrRP+ neural activation in fasted rats that were not shocked retrained before dark chamber re-exposure. Estimation statistics indicated that 12.1% more PrRP+ neurons were activated in shock-retrained, fasted rats pretreated with GRA before dark chamber re-exposure compared to PrRP+ neural activation in similarly shock-retrained, fasted rats treated with saline before dark chamber re-exposure [95%CI: 2.39%, 21.9%].

**Fig. 4.**
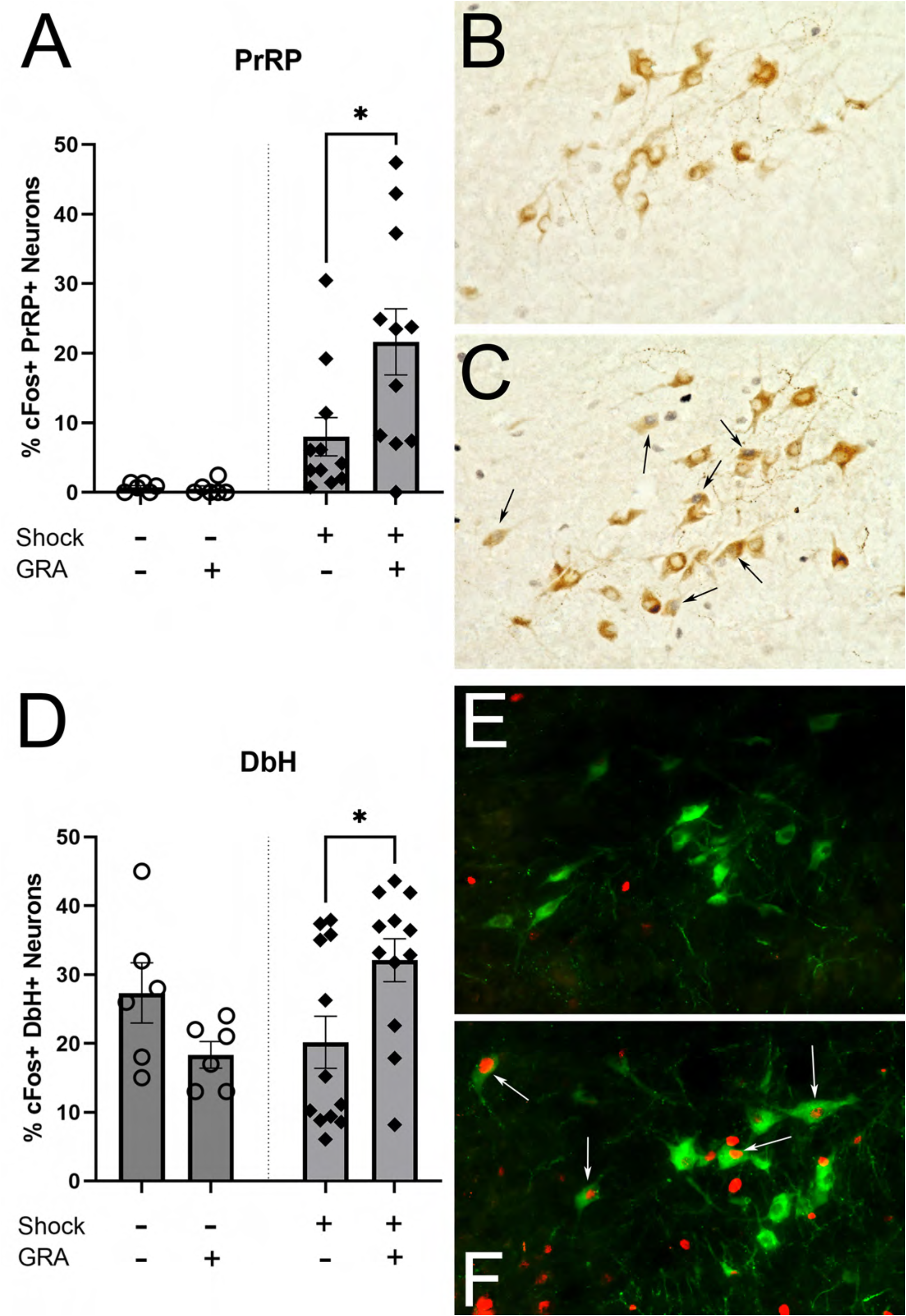
In food deprived male rats, systemic ghrelin receptor antagonist (GRA) administration prior to dark chamber re-exposure increased cFos expression in noradrenergic cNTS neurons in rats that were retrained with footshock (Shock +), but not in rats that were not shock retrained (Shock -). (A) GRA pretreatment increased shock-paired dark chamber-induced activation of the PrRP+ subset of noradrenergic A2 neurons. (B, C) Representative images taken at 40x magnification of nuclear cFos (dark blue) and cytoplasmic PrRP immunolabeling (brown) in saline vehicle-(B) and GRA-treated (C) rats that were shock retrained before being re-exposed to the shock-paired dark chamber. Small black arrows indicate several double-labeled (cFos+) PrRP+ cells. (D) GRA pretreatment also increased shock-paired dark chamber-induced activation of the larger population of DbH+ A2 neurons. (E+F) Representative images taken at 40x magnification of nuclear cFos (red) and cytoplasmic DbH immunolabeling (green) in saline vehicle-(E) and GRA-pretreated (F) rats that were shock retrained before being re-exposed to the shock-paired dark chamber. Small white arrows indicate several double-labeled (cFos+) DbH+ cells. Graphs display group mean +/-SEM, with individual data points plotted. *p<0.05.

Two-way ANOVA also revealed a significant interaction between shock training and GRA treatment [F(1,32) = 7.29; p =0.011] on DbH+ neuron activation (Fig. 4D), but no main effects of either condition. Post-hoc Sidak’s multiple comparisons tests indicated that GRA treatment significantly increased DbH+ neural activation in shock retrained rats after re-exposure to the shock-paired dark chamber (p = 0.024), but no effect of GRA treatment on DbH+ neural activation in non-shocked rats. Estimation statistics indicated that dark chamber re-exposure activated 11.9% more DbH+ neurons in shock-trained rats pretreated with GRA compared to rats treated with saline [95%CI: 1.59%, 20.4%].

Regarding activation of neurons within forebrain regions that receive input from hindbrain A2 neurons, two-way ANOVA indicated a significant main effect of shock training [F(1,30) = 57.08; p < 0.001] and a significant interaction between shock training and GRA treatment [F(1,30) = 5.55; p = 0.025] on dark chamber-induced cFos in the vlBNST (Fig. 5A). Post-hoc Sidak’s multiple comparisons tests indicated significantly more vlBNST cFos activation in GRA-vs. saline-treated rats that had been shock retrained (p = 0.031), but no effect of GRA treatment on vlBNST cFos in non-shocked rats. Estimation statistics indicated that 12.9 more vlBNST neurons per ROI were activated in response to the shock-paired dark chamber in rats pretreated with GRA compared to those treated with saline [95%CI: 3.4, 25.3]. A significant main effect of shock training on PVN cFos also was observed [F(1,32) = 19.54; p < 0.001], with a marginally trending interaction between shock training and GRA treatment [F(1,32) = 2.296; p = 0.14] (Fig. 5D). Post-hoc Sidak’s multiple comparisons tests indicated a trend towards more PVN cFos activation in GRA-vs. saline-treated rats that had been shock retrained before dark chamber re-exposure (p = 0.059), but no effect of GRA treatment on PVN cFos in non-shocked rats. Estimation statistics indicated that 74.6 more PVN neurons/mm^2^ were activated in shock retrained rats pretreated with GRA compared to rats treated with saline [95%CI: 8.64, 137].

**Fig. 5.**
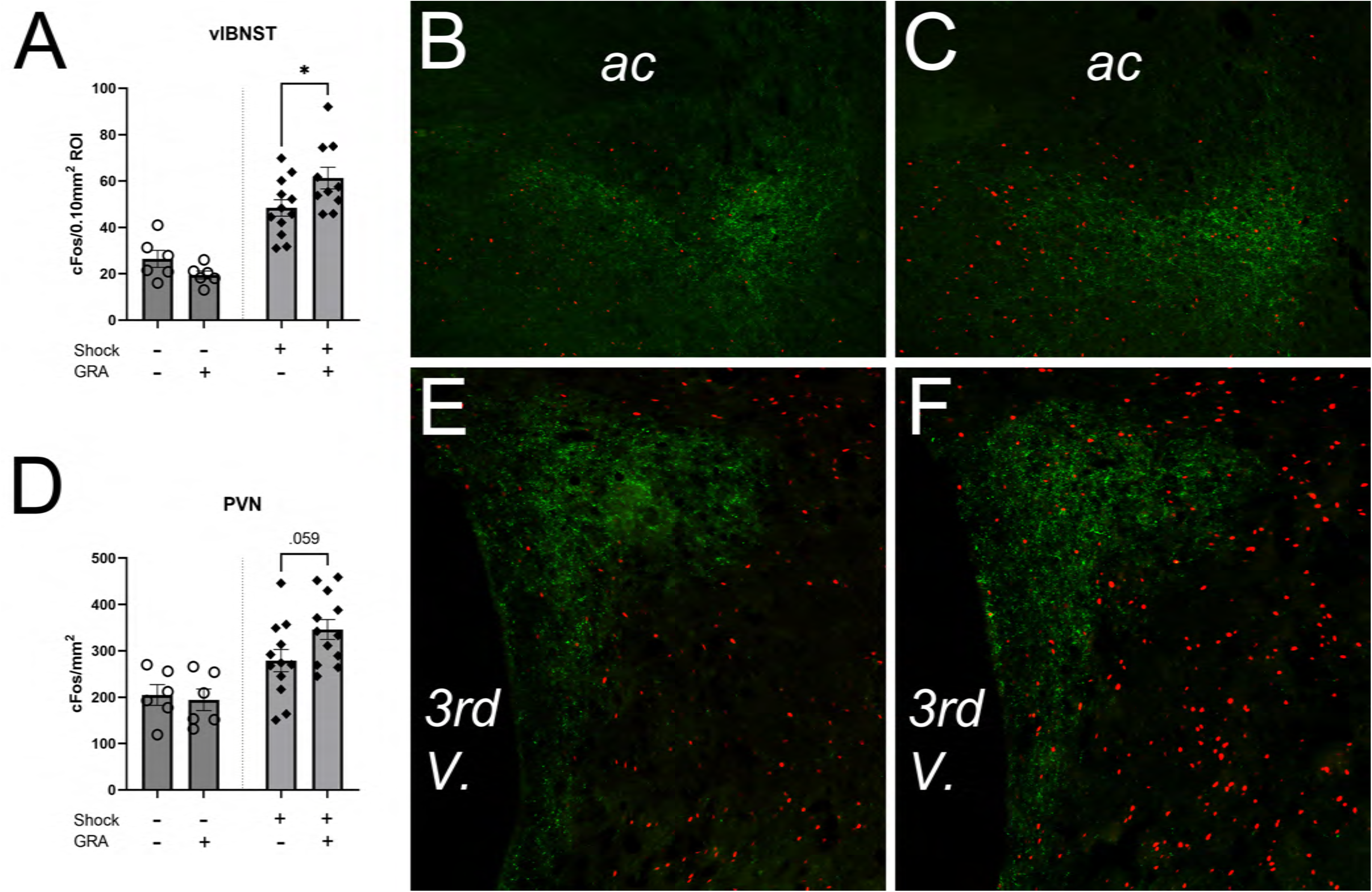
In food deprived male rats that were previously retrained with footshock exposure (Shock +), systemic ghrelin receptor antagonist (GRA) administration increased the ability of dark chamber re-exposure to activate cFos expression in the DbH+ terminal-rich regions of the vlBNST and PVN. GRA did not alter cFos activation in non-shock retrained (Shock -) rats. (A) GRA pretreatment increased shock-paired dark chamber-induced activation of neurons in the DbH+ terminal-rich region of vlBNST. (B+C) Representative images of nuclear cFos (red) and DbH+ terminal labeling (green) in saline vehicle-(B) and GRA-pretreated (C) rats that were re-exposed to the shock-paired dark chamber. ac, anterior commissure. (D) GRA pretreatment tended to increase shock-paired dark chamber-induced activation of neurons in the DbH+ terminal-rich region of the PVN. (E+F) Representative images of nuclear cFos (red) and DbH+ terminal labeling (green) in vehicle-(E) and GRA-pretreated (F) rats that were re-exposed to the shock-paired dark chamber. 3rd V., third ventricle. Graphs display group mean +/-SEM, with individual data points plotted. *p<0.05.

Across all shock retrained rats, Pearson’s r correlations indicated strong positive within-subjects relationships between activation of DbH+ cNTS neurons and activation of neurons in the vlBNST [r(20) = 0.5093; p = 0.0155] (Fig. 6A) and PVN [r(22) = 0.6494; p = 0.0006] (Fig. 6B). Similar positive relationships were observed between activation of PrRP+ cNTS neurons and activation of neurons in the vlBNST [r(20) = 0.5497; p = 0.0080] (Fig. 6C) and PVN [r(22) = 0.5488; p = 0.0055] in shock retrained rats (Fig. 6D). Conversely, there were no significant relationships between cFos expression by PrRP+ or DbH+ neurons within the cNTS and cFos expression in either the PVN or vlBNST in the non-shock retrained rats.

**Fig. 6.**
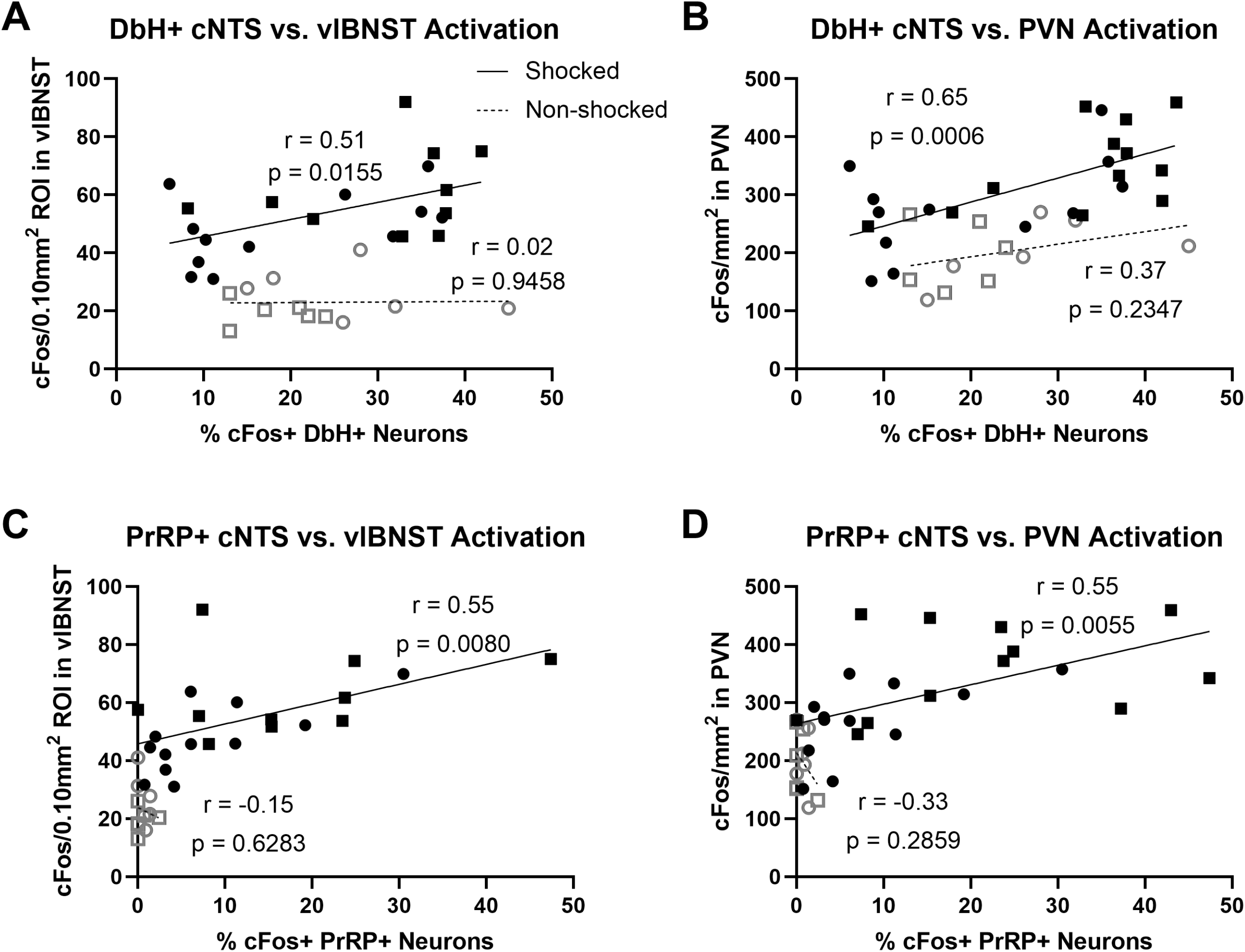
Scatterplots depicting significant positive relationships in shock retrained rats between activation of (A) DbH+ neurons in the cNTS and neurons in the DbH terminal-rich region of the vlBNST, (B) DbH+ neurons and neurons in the DbH terminal-rich region of the PVN, (C) PrRP+ neurons in the cNTS and neurons in the vlBNST, and (D) PrRP+ neurons in the cNTS and neurons in the PVN. Conversely, there were no significant relationships in these neural activation patterns in rats that were not shock retrained before dark chamber re-exposure (Non-shocked).

### Acyl-Ghrelin Levels

One-way ANOVA confirmed a trending effect of food deprivation to increase plasma levels of acyl-ghrelin in adult male rats [F(2,7) = 3.851; p = 0.0745]. Acyl-ghrelin levels in *ad libitum* fed rats averaged 120.1±8.0 pg/ml. Average acyl-ghrelin levels rose to 192.3±24.5 pg/ml following a single overnight fast, and measured 399.9±105.2 pg/ml following a third overnight fast. Thus, there was no decrement of the acyl-ghrelin response to an overnight fast in rats exposed to repeated spaced episodes of overnight fasting.

## Discussion/Conclusion

Results from the present study build on our previous findings that overnight food deprivation reduces conditioned passive avoidance behavior in adult male rats, with concurrent reduction of conditioned context-induced activation of PrRP+ NA neurons in the NTS and downstream target neurons in the vlBNST [7]. Here we demonstrate a role for ghrelin signaling in these fasting-induced behavioral and neural effects. Specifically, we show that pharmacological blockade of fasting-related ghrelin receptor signaling (via systemic administration of GRA) prior to the retention test enhanced conditioned passive avoidance behavior (shown in Fig. 1) and neuronal activation by the aversively conditioned context (Figs. 4, 5). These novel findings support the view that reduced avoidance behavior and reduced neural activation in food-deprived rats is at least partly due to increased ghrelin signaling.

We considered the possibility that reduced passive avoidance behavior in fasted rats with elevated ghrelin is due to a general reduction in memory retrieval, such that GRA may increase passive avoidance by improving memory of the shock-associated context. However, GRA treatment did not improve novel object recognition memory in food deprived rats (shown in Fig. 2). A key distinction between mnemonic processes underlying passive avoidance vs. novel object recognition is that the former but not the latter involves contextual memory [37,38]. Previous research has shown that mice with a global ghrelin receptor knockout and rats with a vagal afferent-specific ghrelin receptor knockdown show impairment in a hippocampal-dependent “novel object in context” task [39,40]. However, GRA effects on passive avoidance memory in the present study were in the opposite direction, such that GRA treatment improved avoidance memory in fasted rats. This discrepancy may be related to chronic rather than acute disruption of ghrelin signaling in the knockout/knockdown studies, which could interfere with other stages of memory formation. Alternatively, reduced performance in the “novel object in context” task by rodents with ghrelin receptor knockout/knockdown could reflect avoidance of the novel object. For example, Davis and colleagues [40] reported a negative shift in discrimination index in their vagal afferent ghrelin receptor knockdown model, such that knockdown rats spent less time exploring the novel object than would be expected if their behavior was due only to impaired memory of the familiar object.

If a fasting-induced elevation in ghrelin signaling is normally anxiolytic, then GRA treatment may reverse this effect to enhance general/innate anxiety-like behavior and avoidance displayed by non-fasted rats. Supporting this idea, previous work demonstrated that overnight food deprivation suppresses innate anxiety-like behavior in the elevated plus maze and light-enhanced acoustic startle tests [6], and in the open field test [41]. In the present study, GRA treatment did not alter innate anxiety-like behavior displayed by food-deprived rats in the open field test (shown in Fig. 3B, C), evidence that ghrelin signaling alone cannot fully explain the anxiolytic effect of food deprivation. A previous study reported that vagal afferent-specific ghrelin receptor knockdown does not alter anxiety-like behavior in the elevated zero maze [40], although rats in that experiment were not food deprived so endogenous ghrelin presumably would have been relatively low in both knockdown and control rats.

Our findings demonstrate that GRA treatment enhanced the ability of re-exposure to the shock-associated context to activate cFos expression in PrRP+ noradrenergic neurons in shock-retrained fasted rats (shown in Fig. 4). These results are consistent with evidence that ghrelin suppresses the activity of vagal sensory afferents [26,27], and suppresses activation of postsynaptic A2 neurons in the cNTS [28]. Our current results also are in line with our previous report that GRA enhances the ability of pharmacological vagal afferent stimulation to activate PrRP+ A2 neurons [20]. In that study [20] and in the current study, GRA treatment did not completely restore A2 neural activation to levels observed in *ad libitum* fed rats (i.e., see [7]). Thus, elevated ghrelin signaling after food deprivation does not fully explain the A2 neural “silencing” effect of negative energy balance. Other contributing factors may include altered mechanosensory signals and/or hormonal signals [6,42,43].

In addition to partial restoration of cNTS neural activation in shock retrained, GRA-treated rats after re-exposure to the shock-paired context, GRA increased shock-paired context-induced activation of neurons within the vlBNST and PVN, two downstream targets of PrRP+ noradrenergic cNTS neurons (shown in Fig. 5). Importantly, GRA treatment prior to dark chamber re-exposure did not influence cFos activation in PrRP+ noradrenergic neurons or within the vlBNST and PVN in rats that were not shocked during training (Figs. 4, 5). These data support the view that ghrelin signaling in fasted rats acts to limit aversive context-induced neural activation, without suppressing neural activation in response to a non-aversive context. Based on significant positive relationships between activation of PrRP+ and DbH+ cNTS neurons and activation of neurons within the vlBNST and PVN (shown in Fig. 6), we hypothesize that activation of A2 cNTS neurons in response to the shock-paired context promotes activation of downstream neurons within the vlBNST and PVN. We previously demonstrated that A2 noradrenergic inputs to the vlBNST and PVN are necessary for stress-induced innate avoidance and behavioral suppression [44–46], and noradrenergic signaling in the BNST is implicated in avoidance behavior conditioned by aversive opiate withdrawal [47,48]. The current results provide new evidence that A2 noradrenergic inputs to the vlBNST and/or PVN also participate in conditioned passive avoidance behavior.

The effect of GRA treatment on behavior and neural activation in the present study could reflect interference with either central or peripheral ghrelin signaling mechanisms. Ghrelin receptor mRNA is expressed by vagal sensory neurons in the nodose ganglia [26,49] and by neurons within the cNTS [50,51], PVN [51], and hippocampus [51,52]. Since systemic administration of the same GRA used in the present study was shown to block the effects of centrally-administered ghrelin [53], central receptor blockade may have contributed to the behavioral and neural effects of GRA in our experiments. Additional work will be needed to differentiate between peripheral/vagally-mediated vs. centrally-mediated ghrelin signaling in conditioned avoidance behavior. Future research also should explore sex differences in passive avoidance behavior and the role of ghrelin signaling (and other interoceptive feedback signals) in that behavior, given our previous report that food deprivation suppresses conditioned passive avoidance in male, but not female, rats [7]. In this regard, recent evidence indicates that female rats have significantly higher plasma levels of acyl-ghrelin compared to male rats under both fed and fasted conditions, and that females may be more sensitive to the anxiolytic effects of fasting-induced ghrelin signaling [25]. We reported that compared to male rats, female rats display markedly less conditioned passive avoidance behavior regardless of metabolic state at testing, consistent with other reports that female rodents show less passive and more active responses to threatening stimuli [7]. Thus, an active avoidance learning and testing approach is likely more suitable to explore potential effects of ghrelin signaling on behavioral (and neural) responses to conditioned aversive stimuli in female rodents.

In summary, our results demonstrate a role for endogenous ghrelin signaling in the ability of negative energy balance to reduce conditioned avoidance behavior in male rats. We further demonstrate that GRA treatment enhances the ability of a footshock-conditioned context to activate PrRP+ A2 noradrenergic neurons, and to activate neurons in the downstream targets of these cNTS neurons (i.e., vlBNST and PVN) in fasted male rats. These findings support the view that interoceptive feedback from body to brain modulates learned avoidance behavior, and that a ghrelin-sensitive circuit contributes to this modulatory effect.

## Statements

### Statement of Ethics

This study protocol was reviewed and approved by Florida State University Animal Care and Use Committee and was conducted in accordance with the National Institutes of Health *Guide for the Care and Use of Laboratory Animals*.

### Author Contributions

CME and LR designed the experiments. CME, IEG, HZ, and TD performed the experiments and analyzed the resulting data. CME, IEG, and LR wrote the manuscript.

### Data Availability Statement

Data are available on request from the corresponding author.

## References

1 American Psychiatric Association. Diagnostic and Statistical Manual of Mental Disorders: DSM 5. Arlington, VA: American Psychiatric Publishing; 2013.

2 Pittig, A, Treanor M, LeBeau RT, Craske MG. The role of associative fear and avoidance learning in anxiety disorders: Gaps and directions for future research. Neurosci Biobehav Rev [Internet]. 2018;88(March):117–40. Available from: https://doi.org/10.1016/j.neubiorev.2018.03.015

3 Dymond S. Overcoming avoidance in anxiety disorders: The contributions of Pavlovian and operant avoidance extinction methods. Neurosci Biobehav Rev [Internet]. 2019;98(October 2018):61–70. Available from: https://doi.org/10.1016/j.neubiorev.2019.01.007

4 Maniscalco JW, Rinaman L. Vagal Interoceptive Modulation of Motivated Behavior. Physiology [Internet]. 2018;33(2):151–67. Available from: http://www.physiology.org/doi/10.1152/physiol.00036.2017

5 Lutter M, Sakata I, Osborne-Lawrence S, Rovinsky SA, Anderson JG, Jung S, et al. The orexigenic hormone ghrelin defends against depressive symptoms of chronic stress. Nat Neurosci [Internet]. 2008 Jul 15;11(7):752–3. Available from: http://www.nature.com/articles/nn.2139

6 Maniscalco JW, Zheng H, Gordon PJ, Rinaman L. Negative Energy Balance Blocks Neural and Behavioral Responses to Acute Stress by “Silencing” Central Glucagon-Like Peptide 1 Signaling in Rats. J Neurosci [Internet]. 2015 Jul 29;35(30):10701–14. Available from: https://www.jneurosci.org/lookup/doi/10.1523/JNEUROSCI.3464-14.2015

7 Edwards CM, Dolezel T, Rinaman L. Sex and metabolic state interact to influence expression of passive avoidance memory in rats: Potential contribution of A2 noradrenergic neurons. Physiol Behav [Internet]. 2021;239:113511. Available from: https://doi.org/10.1016/j.physbeh.2021.113511

8 Klarer M, Arnold M, Gunther L, Winter C, Langhans W, Meyer U. Gut Vagal Afferents Differentially Modulate Innate Anxiety and Learned Fear. J Neurosci [Internet]. 2014;34(21):7067–76. Available from: http://www.jneurosci.org/cgi/doi/10.1523/JNEUROSCI.0252-14.2014

9 Foster JA, Rinaman L, Cryan JF. Stress & the gut-brain axis: Regulation by the microbiome. Neurobiol Stress [Internet]. 2017;7:124–36. Available from: https://doi.org/10.1016/j.ynstr.2017.03.001

10 Holt MK, Rinaman L. The role of nucleus of the solitary tract glucagon-like peptide-1 and prolactin-releasing peptide neurons in stress: anatomy, physiology and cellular interactions. Br J Pharmacol. 2021;(January):1–17.

11 Shapiro RE, Miselis RR. The central organization of the vagus nerve innervating the stomach of the rat. J Comp Neurol. 1985;238:473–88.

12 Altschuler SM, Bao X, Bieger D, Hopkins DA, Miselis RR. Viscerotopic representation of the upper alimentary tract in the rat: Sensory ganglia and nuclei of the solitary and spinal trigeminal tracts. J Comp Neurol. 1989;283(2):248–68.

13 Raybould HE. Gut chemosensing: Interactions between gut endocrine cells and visceral afferents. Auton Neurosci [Internet]. 2010 Feb;153(1–2):41–6. Available from: https://linkinghub.elsevier.com/retrieve/pii/S1566070209004111

14 Mayer EA. Gut feelings: The emerging biology of gut-”brain communication. Nat Rev Neurosci. 2011;12(8):453–66.

15 Dockray GJ. Gastrointestinal hormones and the dialogue between gut and brain. J Physiol. 2014;592(14):2927–41.

16 Appleyard SM, Marks D, Kobayashi K, Okano H, Low MJ, Andresen MC. Visceral afferents directly activate catecholamine neurons in the solitary tract nucleus. J Neurosci. 2007;27(48):13292–302.

17 Rinaman L. Hindbrain noradrenergic A2 neurons: Diverse roles in autonomic, endocrine, cognitive, and behavioral functions. Am J Physiol - Regul Integr Comp Physiol. 2011;300(2):222–35.

18 Mönnikes H, Lauer G, Arnold R. Peripheral administration of cholecystokinin activates c-fos expression in the locus coeruleus/subcoeruleus nucleus, dorsal vagal complex and paraventricular nucleus via capsaicin-sensitive vagal afferents and CCK-A receptors in the rat. Brain Res. 1997;770(1–2):277–88.

19 Maniscalco JW, Rinaman L. Overnight food deprivation markedly attenuates hindbrain noradrenergic, glucagon-like peptide-1, and hypothalamic neural responses to exogenous cholecystokinin in male rats. Physiol Behav [Internet]. 2013 Sep;121:35–42. Available from: https://www.ncbi.nlm.nih.gov/pmc/articles/PMC3624763/pdf/nihms412728.pdf

20 Maniscalco JW, Edwards CM, Rinaman L. Ghrelin signaling contributes to fasting-induced attenuation of hindbrain neural activation and hypophagic responses to systemic cholecystokinin in rats. Am J Physiol - Regul Integr Comp Physiol. 2020;318(5):R1014–23.

21 Zhu L, Onaka T. Involvement of medullary A2 noradrenergic neurons in the activation of oxytocin neurons after conditioned fear stimuli. Eur J Neurosci. 2002;16(11):2186–98.

22 Morales T, Sawchenko PE. Brainstem prolactin-releasing peptide neurons are sensitive to stress and lactation. Neuroscience. 2003;121(3):771–8.

23 Tschop M, Smiley DL, Heiman ML. Ghrelin induces adiposity in rodents. Nature. 2000;407(6806):908–13.

24 Cummings DE, Purnell JQ, Frayo RS, Schmidova K, Wisse BE, Weigle DS. A preprandial rise in plasma ghrelin levels suggests a role in meal inititation in humans. Diabetes. 2001;50:1714–9.

25 Börchers S, Krieger J-P, Maric I, Carl J, Abraham M, Longo F, et al. From an Empty Stomach to Anxiolysis: Molecular and Behavioral Assessment of Sex Differences in the Ghrelin Axis of Rats. Front Endocrinol (Lausanne). 2022;13(June):1–16.

26 Date Y, Murakami N, Toshinai K, Matsukura S, Niijima A, Matsuo H, et al. The role of the gastric afferent vagal nerve in Ghrelin-induced feeding and growth hormone secretion in rats. Gastroenterology. 2002;123(4):1120–8.

27 Date Y, Toshinai K, Koda S, Miyazato M, Shimbara T, Tsuruta T, et al. Peripheral interaction of ghrelin with cholecystokinin on feeding regulation. Endocrinology. 2005;146(8):3518–25.

28 Cui RJ, Li X, Appleyard SM. Ghrelin inhibits visceral afferent activation of catecholamine neurons in the solitary tract nucleus. J Neurosci. 2011;31(October 2009):3484–92.

29 Spencer SJ, Xu L, Clarke MA, Lemus M, Reichenbach A, Geenen B, et al. Ghrelin regulates the hypothalamic-pituitary-adrenal axis and restricts anxiety after acute stress. Biol Psychiatry. 2012;72(6):457–65.

30 Harmatz ES, Stone L, Lim SH, Lee G, Mcgrath A, Peng X, et al. Central ghrelin resistance permits the overconsolidation of fear memory. Biol Psychiatry. 2018;81(12):1003–13.

31 Watson RE, Wiegand SJ, Clough RW, Hoffman GE. Use of cryoprotectant to maintain long-term peptide immunoreactivity and tissue morphology. Peptides. 1986;7(1):155–9.

32 Maruyama M, Matsumoto H, Fujiwara K, Kitada C, Hinuma S, Onda H, et al. Immunocytochemical localization of prolactin-releasing peptide in the rat brain. Endocrinology. 1999;140(5):2326–33.

33 Lin SHS, Leslie FM, Civelli O. Neurochemical properties of the prolactin releasing peptide (PrRP) receptor expressing neurons: Evidence for a role of PrRP as a regulator of stress and nociception. Brain Res. 2002;952(1):15–30.

34 Ruxton GD. The unequal variance t-test is an underused alternative to Student’s t-test and the Mann-Whitney U test. Behav Ecol. 2006;17(4):688–90.

35 Calin-Jageman R, Cumming G. Estimation for Better Inference in Neuroscience. eNeuro. 2019;6(August):1–11.

36 Ho J, Tumkaya T, Aryal S, Choi H, Claridge-Chang A. Moving beyond P values: data analysis with estimation graphics. Nat Methods [Internet]. 2019;16(7):565–6. Available from: http://dx.doi.org/10.1038/s41592-019-0470-3

37 Atucha E, Roozendaal B. The inhibitory avoidance discrimination task to investigate accuracy of memory. Front Behav Neurosci. 2015;9(March):1–8.

38 Lueptow LM. Novel object recognition test for the investigation of learning and memory in mice. J Vis Exp. 2017;2017(126):1–9.

39 Diano S, Farr SA, Benoit SC, McNay EC, Da Silva I, Horvath B, et al. Ghrelin controls hippocampal spine synapse density and memory performance. Nat Neurosci. 2006;9(3):381–8.

40 Davis EA, Wald HS, Suarez AN, Zubcevic J, Liu CM, Cortella AM, et al. Ghrelin Signaling Affects Feeding Behavior, Metabolism, and Memory through the Vagus Nerve. Curr Biol [Internet]. 2020;30(22):4510-4518.e6. Available from: https://doi.org/10.1016/j.cub.2020.08.069

41 Heiderstadt KM, McLaughlin RM, Wright DC, Walker SE, Gomez-Sanchez CE. The effect of chronic food and water restriction on open-field behaviour and serum corticosterone levels in rats. Lab Anim. 2000;34(1):20–8.

42 Dallman MF, Akana SF, Bhatnagar S, Bell ME, Choi SJ, Chu A, et al. Starvation: Early signals, sensors, and sequelae. Endocrinology. 1999;140(9):4015–23.

43 Maniscalco JW, Rinaman L. Interoceptive modulation of neuroendocrine, emotional, and hypophagic responses to stress. Physiol Behav [Internet]. 2017 Jul;176:195–206. Available from: https://linkinghub.elsevier.com/retrieve/pii/S0031938416311362

44 Banihashemi L, Rinaman L. Noradrenergic inputs to the bed nucleus of the stria terminalis and paraventricular nucleus of the hypothalamus underlie hypothalamic-pituitary-adrenal axis but not hypophagic or conditioned avoidance responses to systemic yohimbine. J Neurosci. 2006;26(44):11442–53.

45 Rinaman L, Dzmura V. Experimental dissociation of neural circuits underlying conditioned avoidance and hypophagic responses to lithium chloride. Am J Physiol - Regul Integr Comp Physiol. 2007;293(4):1495–503.

46 Zheng H, Rinaman L. Yohimbine anxiogenesis in the elevated plus maze requires hindbrain noradrenergic neurons that target the anterior ventrolateral bed nucleus of the stria terminalis. Eur J Neurosci [Internet]. 2013 Apr;37(8):1340–9. Available from: https://www.ncbi.nlm.nih.gov/pmc/articles/PMC3624763/pdf/nihms412728.pdf

47 Aston-Jones G, Delfs JM, Druhan J, Zhu Y. The bed nucleus of the stria terminalis. A target site for noradrenergic actions in opiate withdrawal. Ann N Y Acad Sci. 1999;877(151):486–98.

48 Delfs JM, Zhu Y, Druhan JP, Aston-Jones G. Noradrenaline in the ventral forebrain is critical for opiate withdrawal-induced aversion. 2000;403(January).

49 Burdyga G, Varro A, Dimaline R, Thompson DG, Dockray GJ. Ghrelin receptors in rat and human nodose ganglia: Putative role in regulating CB-1 and MCH receptor abundance. Am J Physiol - Gastrointest Liver Physiol. 2006;290(6):1289–97.

50 Lin Y, Matsumura K, Fukuhara M, Kagiyama S, Fujii K, Iida M. Ghrelin Acts at the Nucleus of the Solitary Tract to Decrease Arterial Pressure in Rats. Hypertension. 2004;43(5):977–82.

51 Zigman JM, Jones JE, Lee CE, Saper CB, Elmquist JK. Expression of ghrelin receptor mRNA in the rat and the mouse brain. J Comp Neurol. 2006;494(3):528–48.

52 Guan XM, Yu H, Palyha OC, McKee KK, Feighner SD, Sirinathsinghji DJS, et al. Distribution of mRNA encoding the growth hormone secretagogue receptor in brain and peripheral tissues. Mol Brain Res. 1997;48(1):23–9.

53 Brockway ET, Krater KR, Selva JA, Wauson SER, Currie PJ. Impact of [d-Lys3]-GHRP-6 and feeding status on hypothalamic ghrelin-induced stress activation. Peptides [Internet]. 2016;79:95–102. Available from: http://dx.doi.org/10.1016/j.peptides.2016.03.013

